# Neural noise is associated with age-related neural dedifferentiation

**DOI:** 10.1101/2022.11.17.516990

**Authors:** Rachelle E. Pichot, Daniel J. Henreckson, Morgan Foley, Joshua D. Koen

## Abstract

Age-related neural dedifferentiation – reductions in the selectivity and precision of neural representations – contributes to cognitive aging and is thought to result from age increases in neural noise. This research has primarily used fMRI to examine age-related reductions in neural selectivity for different categories of visual stimuli. The present experiment used EEG to examine the link between neural noise and age-related neural dedifferentiation indexed by the scene-selective (P200) and face-selective (N170) ERP components. Young and older adults viewed images of scenes, objects, and faces during a 1-back task. Whereas both the P200 and N170 showed age-related slowing of peak latency, only the P200 showed age-related reductions in amplitude that were independent of visual and contrast acuity. We also examined the relationship between the ERP peak measures and an index of neural noise, namely the 1/f exponent of the frequency power spectrum. For the P200 amplitude, higher levels of neural noise were associated with smaller P200 amplitudes in young, but not older adults. In contrast, there was an age-invariant relationship between neural noise and N170 amplitude in the left hemisphere with higher levels of neural noise being associated reduced N170 amplitudes. While the present findings provide novel empirical evidence broadly consistent with predictions from computational models of neural dedifferentiation, the results also highlight potential limitations of the computational model that necessitate revision. The results also suggest that, at least for the P200, maintaining levels of neural noise similar to young adults might preserve levels of neural selectivity.

**Significance Statement:** A prominent theory of cognitive aging proposes that age-related cognitive decline results from increases in neural noise that reduce the selectivity of neural representations. We examined this predicted link between neural selectivity and neural noise with ERP components that show selectivity for scenes (P200) and faces (N170) and the 1/f aperiodic exponent measure of neural noise. The amplitude for the scene-selective, but not face-selective, ERP component was reduced in older adults, with both components showing age-related slowing. Critically, older adults with higher levels of neural noise showed lower levels of neural selectivity for scenes, but not faces. While these results provide some evidence supporting computational models of neural dedifferentiation, they also highlight important limitations of the model that require revision.

## Introduction

Healthy aging is associated with declines in many cognitive abilities (Harada et al., 2013; Salthouse, 2010, 2019). An influential theory of cognitive aging proposes that age-related neural dedifferentiation – reductions in the distinctiveness or precision of neural representations – contributes to age-related cognitive decline (Koen & Rugg, 2019; Koen et al., 2020; Li et al., 2001; Li & Rieckmann, 2014a). Computational models of neural dedifferentiation propose that age-related reductions in neuromodulatory drive increases neural noise (i.e., decreases the signal-to-noise ratio of neural activity; Li, 2013; Li & Rieckmann, 2014a; Mather & Harley, 2016; Seaman et al., 2019). Increases in neural noise, in turn, contribute to reductions in the selectivity or precision of neural representations that negatively affect cognition. This model predicts that neural selectivity will decrease with age showing an age-invariant relationship with measures of cognition (Koen et al., 2020; Koen & Rugg, 2019). The majority of findings support both predictions with older adults showing reduced neural selectivity in visual, auditory, and motor stimuli (Carp et al., 2011; Chamberlain et al., 2021; Koen, 2022; Koen et al., 2019; Lalwani et al., 2019; D. C. Park et al., 2004; J. Park et al., 2012; Simmonite & Polk, 2022; Srokova et al., 2020; Voss et al., 2008) and age-invariant associations between neural selectivity and measures of cognitive performance (Berron et al., 2018; Bowman et al., 2019; Du et al., 2016; Koen, 2022; Koen et al., 2019; J. Park et al., 2010; Srokova et al., 2020; Yassa et al., 2011). However, relatively little work has been done to examine the critical prediction from computational models of neural dedifferentiation that neural noise contributes to age-related reductions in neural selectivity.

Age-related increases in neural noise can be measured as increases in spontaneous (i.e., task-unrelated) or asynchronous neural activity (Li et al., 2001; Voytek & Knight, 2015; Li & Sikström, 2002; Hong & Rebec, 2012). Findings from single-unit recordings in non-human animals have provided the strongest evidence linking age-related increases in spontaneous neural activity (i.e., neural noise) to decreases in neural selectivity (Ding et al., 2017; Engle & Recanzone, 2013; Hua et al., 2006; Juarez-Salinas et al., 2010; Leventhal et al., 2003; Liang et al., 2010; Schmolesky et al., 2000; Yang et al., 2008; Yu et al., 2006; Zhang et al., 2008; but see Costa et al., 2016; Turner et al., 2005). In studies involving humans, age-related increases in neural noise are primarily observed using measures of population level neural activity derived from examining the 1/f aperiodic component of the EEG power spectrum (Dave et al., 2018; Donoghue et al., 2020; Voytek et al., 2015; Voytek & Knight, 2015). The 1/f component of the power spectrum reflects aperiodic neural activity and is associated with excitatory-inhibitory balance in neural activity, which plays a critical role in neural noise (Freeman & Zhai, 2009; R. Gao, 2015). As noted previously, it remains unclear whether measures of neural noise derived from EEG are related to neural selectivity in young and older adult humans.

The purpose of the present study is to test the critical prediction from computational models of neural dedifferentiation that age-related increases in neural noise contribute to age-related reductions in neural selectivity (Li et al., 2001; Li & Rieckmann, 2014a). In the present experiment, EEG was recorded while young and older adults completed a 1-back task with scene, object, and face images (see Figure 1A). Measures of neural selectivity were examined by measuring two ERP components: a scene-selective P200 (Harel et al., 2016) and the face-selective N170 (Daniel & Bentin, 2012; R. Gao, 2015; Rousselet et al., 2007; for review, see Rossion & Jacques, 2012). In addition to measuring the amplitude of these two components, we measured the latency and onset of the P200 and N170. We related these ERP peak measures with the 1/f exponent measure of neural noise derived from the EEG frequency power spectrum (Donoghue et al., 2020).

**Figure 1.**
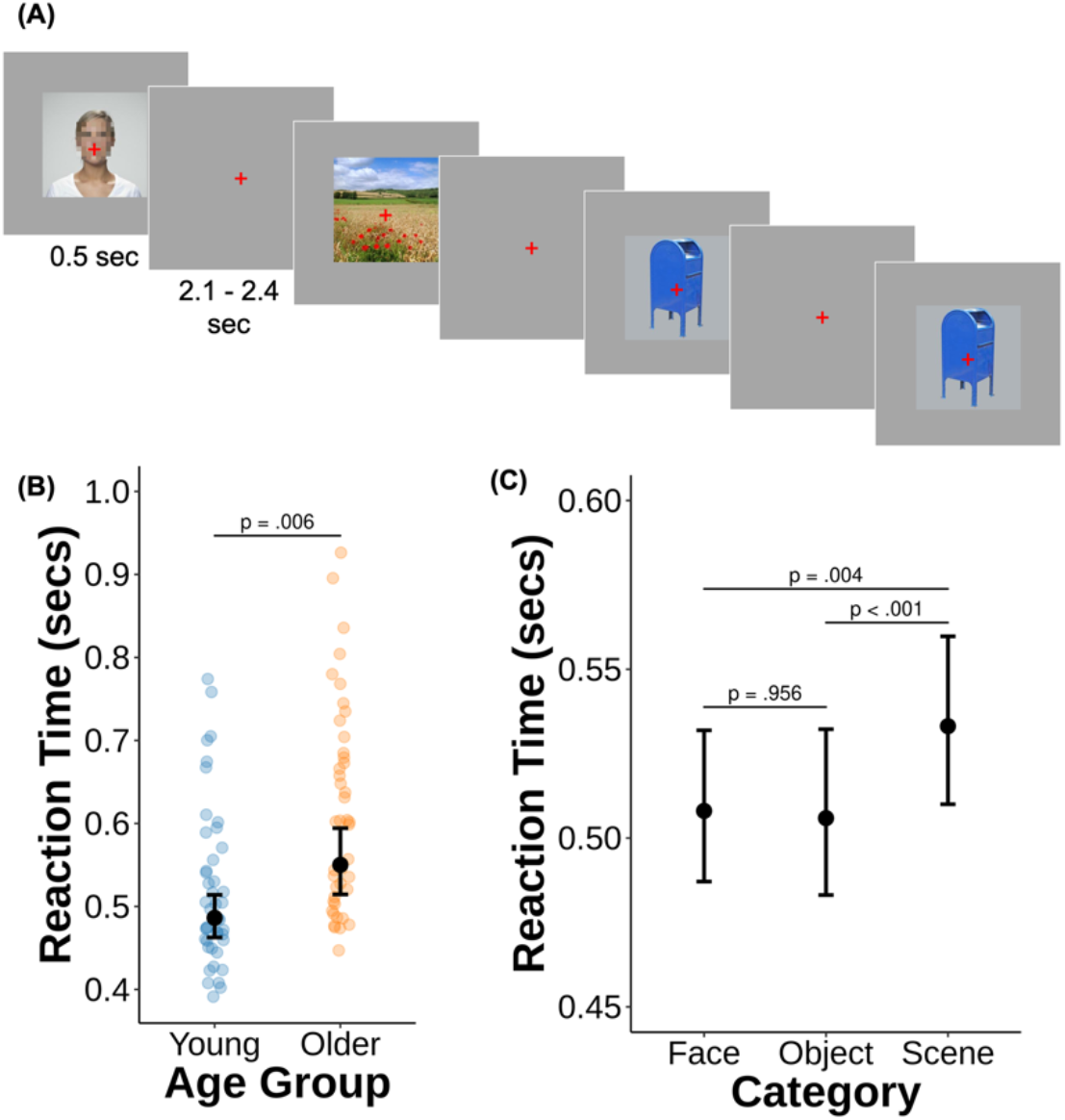
Depiction of the experimental paradigm and behavioral results. (A) Schematic overview of the scene, object, and face 1-back task that participants completed while undergoing EEG recording. Participants were instructed to press a button with their right index finger if an image repeated itself. (B) The analysis of reaction times (RTs) for correct 1-back judgments demonstrated a significant effect of age, whereby older adults were slower to respond relative to young adults. (C) There was also a significant effect of image category on RTs during correct 1-back judgments. Correct detection of repeated scenes was slower than for both faces and objects, and there was no significant difference in RTs between faces and objects. The yellow and orange points depict individual participant RTs. Black points reflect the model implied mean RT with error bars representing the 95% confidence intervals of the estimate.

## Methods

### Ethics Statement

The institutional review board of the University of Notre Dame approved the experimental procedures described below. All participants provided written informed consent before participating in each experimental session.

### Participants

The sample contributing data to the current report comprised 44 young (18-30 years of age) and 44 older (65-82 years of age) cognitively normal adults (see Table 1 for demographics and scores on the neuropsychological test battery). Participants were recruited from the University of Notre Dame and the surrounding areas and compensated for time at $15/hour and travel expenses. All participants were native English speakers, right-handed, had normal or correct-to-normal vision, had no self-reported history of substance abuse, psychological or neurological issues, seizure, stoke, neuroactive medications, severe head injury or diagnosed concussion within the past 3 months, memory disorder, heart disease or arrhythmia, uncontrolled hypertension, diabetes, COVID-19 risk factors (i.e., liver disease, immunocompromised, moderate to severe asthma; except being over the age of 65 in the case of older adults) and performed within the norms of a neuropsychological test battery. Data from an additional three participants were collected and excluded from analyses due to having greater than 25% of EEG epochs rejected (1 young adult and 1 older adult) or due to a misunderstanding of the task instructions (1 older adult).

**Table 1.** Sample demographics and neuropsychological test scores.

|  | Young | Older | *t*(86) | *p* | Cohen's<br>*d* |
| --- | --- | --- | --- | --- | --- |
| N | 44 | 44 |  |  |  |
| Age | 20.61<br>(2.73) | 70.75<br>(4.58) |  |  |  |
| Sex/Gender (F/M) | 25/19 | 25/19 |  |  |  |
| Ethnicity |  |  |  |  |  |
| Hispanic or Latino | 5 | 0 |  |  |  |
| Not Hispanic or Latino | 39 | 43 |  |  |  |
| Not reported/Unknown | 0 | 1 |  |  |  |
| Race |  |  |  |  |  |
| White | 34 | 43 |  |  |  |
| Asian | 4 | 0 |  |  |  |
| More than one race | 6 | 0 |  |  |  |

| Not reported/Unknown | 0 | 1 |  |  |  |
| --- | --- | --- | --- | --- | --- |
| Education (years) | 13.80<br>(1.80) | 17.05<br>(2.61) | 6.81 | < .001 | 1.47 |
| MOCA | 28.75<br>(1.37) | 26.09<br>(1.67) | 8.18 | < .001 | 1.76 |
| CVLT Short Delay Free Recall | 13.32<br>(2.44) | 9.57<br>(2.56) | 7.03 | < .001 | 1.52 |
| CVLT Long Delay Free Recall | 13.57<br>(2.21) | 9.95<br>(2.87) | 6.61 | < .001 | 1.43 |
| CVLT Recognition Hits | 15.25<br>(1.06) | 14.66<br>(1.33) | 2.31 | .023 | 0.50 |
| CVLT Recognition False Alarms | 0.45<br>(0.82) | 2.59<br>(3.27) | 4.21 | < .001 | 0.91 |
| Logical Memory I | 29.32<br>(6.56) | 26.00<br>(4.72) | 2.72 | .008 | 0.59 |
| Logical Memory II | 25.84<br>(6.91) | 21.93<br>(5.61) | 2.91 | .005 | 0.63 |
| SDMT Written | 68.45<br>(13.45) | 48.64<br>(8.13) | 8.36 | < .001 | 1.80 |
| SDMT Oral | 83.23<br>(16.17) | 57.20<br>(8.88) | 9.36 | < .001 | 2.02 |
| Digit Span | 20.20<br>(4.56) | 17.00<br>(3.86) | 3.56 | .001 | 0.77 |
| Trails A (s) | 17.98<br>(4.31) | 28.97<br>(10.43) | 6.46 | < .001 | 1.39 |
| Trails B (s) | 41.69<br>(14.82) | 64.26<br>(22.40) | 5.57 | < .001 | 1.20 |
| Categories (Animals) | 27.43<br>(4.77) | 22.93<br>(6.25) | 3.80 | < .001 | 0.82 |
| Stroop Interference | 1.50<br>(0.30) | 2.00<br>(0.49) | 5.69 | < .001 | 1.23 |
| WASI – Matrix Reasoning | 23.41<br>(2.44) | 19.27<br>(4.07) | 5.78 | < .001 | 1.25 |
| WASI – Vocabulary | 43.07<br>(3.17) | 42.73<br>(5.58) | 0.35 | .725 | 0.08 |
| TOPF | 51.41<br>(9.35) | 51.16<br>(10.61) | 0.12 | .907 | 0.03 |
| Visual Acuity (logMAR) | -0.14<br>(0.08) | -0.01<br>(0.08) | 7.39 | < .001 | 1.59 |
| Contrast Acuity (logCS) | 1.36<br>(0.24) | 0.83<br>(0.27) | 9.80 | < .001 | 2.11 |

*Note.* Standard deviations are provided in parentheses with the mean values. Statistical differences between groups determined by independent samples *t*-test assuming equal variances.
MOCA: Montreal Cognitive Assessment, CVLT: California Verbal Learning Test, SDMT: Symbol Digit Modalities Test, Logical Memory: from Wechsler Memory Scale, WASI: Wechsler Abbreviated Scale of Intelligence, TOPF: Test of Premorbid Functioning.

### Neuropsychological Test Battery

All participants completed a neuropsychological test battery that included the Montreal Cognitive Assessment (MoCA; Nasreddine et al., 2005), the California Verbal Learning Test, Third Edition (Delis et al., 2017), the written and oral forms of the Symbol Digit Modalities Test (Smith, 1982), the Digit Span from the Wechsler Adult Intelligence Scale-Revised (Wechsler, 1981), Trail Making Test parts A and B (Reitan & Wolfson, 1985), Logical Memory I and II from the Wechsler Memory Scale – Revised (Wechsler, 2009), the Vocabulary and Matrix Reasoning portions of the Wechsler Abbreviated Scale of Intelligence (Wechsler, 2011), the Test of Premorbid Functioning (Pearson, 2009), Category Fluency test for animals (Rosen, 1980; Tombaugh et al., 1999), and the Victoria Stroop Test (Regard, 1981; Troyer et al., 2006). Participants were excluded prior to the EEG experiment if they (1) scored below a 23 on the MoCA (Carson et al., 2018), (2) scored more than 1.5 standard deviations under the age- and education-normed mean performance on any one memory measure (i.e., CVLT short or long delay cued or free recall, CVLT recognition discriminability, and Logical Memory I and II), or (3) scored more than 1.5 standard deviations below age-normed and, if available education-normed, mean performance for two or more of the other measures in the battery. These exclusionary criteria for the neuropsychological test battery were to minimize the possibility that older adults were in the early stages of mild cognitive impairment. Additionally, participants’ visual acuity and contrast sensitivity was measured using the Super Vision Test (Rabin et al., 2009). Neither visual acuity nor contrast sensitivity were used as exclusionary criteria.

### Experimental Task

#### Materials

The stimuli for this task comprised 96 object images from the BOSS database (Brodeur, 2014), 96 scene images from the Konkle et al. (2012) database, and 96 faces from the London Face Database (DeBruine & Jones, 2021). Half of the objects and scenes depicted manmade images (e.g., tools, urban landscapes) and the other half depicted natural images (e.g., vegetables, rural scenes). All scene images were of outdoor locations and did not depict animals or people. Faces included males and females from multiple races and ethnicities with neutral expressions.

The images were used to create yoked stimulus sets that were each presented to one young and one older adult. Each yoked stimulus set comprised 2 lists, each with 144 novel (i.e., first) presentation trials and 18 repeated trials for a total of 288 novel presentation trials and 36 repeated trials. A total of 12.5% of the images were randomly assigned to be repeated as 1-back trials, resulting in 11.1% of all trials being repeated images. There were an equal number of each image sub-category (e.g., natural vs. manmade) within each condition (novel vs. repeated) in each block. The stimulus lists were pseudorandomized such that there were no more than 3 repetitions of the same image category and no more than two consecutive images labeled 1-back trials. The pseudorandomization was done prior to inserting the 1-back trials in the trial sequence.

Presentation of all tasks was controlled with *PsychoPy* (Peirce, 2007; Peirce et al., 2019) on a Windows 10 PC computer with a BenQ XL4340 monitor (100Hz frame refresh rate) using frame-rate timing. Participants were seated approximately 57 cm from the monitor for all tasks. Images were presented on the computer screen subtending a visual angle of approximately 10.0° x 10.0° centered on a grey background. Object images were overlaid on a light grey background of the same size as the scene and face images to approximately equate the area of the monitor taken up by the object, face, and scene images. A red fixation cross (1° letter height) always remained at the center of the screen.

#### Procedure

The experiment took place over two sessions conducted on separate days. The first session consisted of the neuropsychological test battery and the second session consisted of the experimental EEG tasks. In the second session, participants completed the following EEG tasks in order: (1) a 1-back task with scene, object, and face stimuli, (2) a C1 wave and oddball task (cf. Kappenman & Luck, 2012), and a (3) recognition memory task (cf. Koen et al., 2019). The present report focuses on the EEG/ERP data from the 1-back task. Data from the other two tasks will be the focus of other reports.

The procedure for the 1-back task was modeled after the paradigm used by Harel and colleagues (2016; see Figure 1A). Participants were instructed to pay attention to each image and, when they saw an image repeat, to press a button on a response pad (LabHackers, Inc.) with their right index finger as quickly as possible. Participants were instructed that if they did not see the image repeat immediately, the image would not repeat again in the task. Each image was presented for 0.5 secs and followed by an inter-trial interval between 1.6 secs and 1.9 secs (in 0.01 secs intervals). This resulted in a stimulus-onset asynchrony range of 2.1 to 2.4 secs. Participants were instructed to keep their eyes fixated on the red fixation cross in the center of the screen through the task. Participants were given a break halfway throughout the task (i.e., after 162 trials). A brief practice task was completed prior to the critical phase to ensure participants understood the instructions.

### Dependent Measures from the Behavioral Data

The dependent measures from the 1-back task included the proportion of accurate no responses to first presentation trials, the proportion of accurate button presses to 1-back trials, and reaction times (RTs, in secs) for accurate 1-back judgments. Accuracies were near ceiling (see Results) and are provided for descriptive purposes only. Statistical analyses were carried out on RTs to examine differences in RTs as a function of age and image category.

### EEG Acquisition and Preprocessing

Continuous EEG was recorded with an actiCHamp amplifier (Brain Products, Munic, Germany) from 64 Ag/AgCI active electrodes (Afz, Fz, FCz, Cz, CPz, Pz, Poz, Oz, Fp1/2, AF3/4/7/8, F1/2/3/4/5/6/7/8, FC1/2/3/4/5/6/7/8, C1/2/3/4/5/6, T7/8, CP1/2/3/4/5/6, TP7/8/9/10, P1/2/3/4/5/6/7/8, PO3/4/7/8, and O1/2) mounted in an actiCAP (Brain Products; Munich, Germany) according to the extended 10-20 system (Nuwer et al., 1998). Data were referenced online to FCz with the ground located at Fpz. Electrode impedances were lowered at or below 10 kΩ before data collection began. Impedances were checked and adjusted, if needed, before each of the three tasks in the experiment. EEG was recorded at DC and digitized at 1000 Hz. The BIO2AUX adaptor was used to acquire and monitor vertical and horizontal electrooculography (EOG). Vertical EOG was monitored with two passive electrodes placed above and below the right eye, with a ground electrode on the right check. Horizontal EOG was monitored with two electrodes placed on the outer canthi of each eye, with a ground electrode placed on the left cheek. Impedance was not monitored for the EOG electrodes. The EOG recordings were primarily used for online monitoring during EEG recording and during preprocessing for identification of blinks occurring around stimulus onset.

Offline analysis of the EEG data was performed using *mne-Python* (version 0.23.4; Gramfort et al., 2013). First, the photosensor data were used to adjust event onset latencies (i.e., markers) using an in-house algorithm. The algorithm first determined the threshold value of an event as 85% of the maximum photosensor value. Then, in each segment surrounding an event marker, the algorithm found the first sample that exceeded the threshold value and marked that as the onset value. An event latency was adjusted if the delay between the original marker and the photosensor-derived onset value was delayed more than half the frame rate (50 Hz or 5 ms).

Next, the continuous data were resampled to 250 Hz and high-pass filtered at 0.10 Hz with a finite-impulse response filter (zero-phase shift; -6 dB cutoff = 0.05 Hz; transition bandwidth = 0.10 Hz). The data were then epoched from -1000 ms to 1000 ms locked to stimulus onset for the 1-back task. Epochs with voltages exceeding ±150 µV, peak-to-peak values exceeding threshold determined with the *get_rejection_threshold* function implemented in *autoreject* (Jas et al., 2017) on 6 or more channels, and blinks at stimulus onset (150 µV peak-to-peak value threshold in a ±100 ms window) in the VEOG channel were automatically flagged for visual inspection and then rejected if determined to contain an artifact. Bad channels were identified manually and using the Random Sample Consensus algorithm (Bigdely-Shamlo et al., 2015; Fischler & Bolles, 1981) as implemented in *autoreject*. Next, artifact correction was performed using independent components analysis (ICA; Jung et al., 2000) using the Piccard algorithm (Ablin et al., 2018). Note that ICA was performed on a duplicate EEG dataset that was processed in the same manner described above, with the exception that the continuous data were first filtered at 1 Hz with a finite-impulse response filter (zero-phase shift; -6 dB cutoff = 0.5 Hz; transition bandwidth = 0.50 Hz). Ocular and other artifactual components (e.g., muscle, heart) were visually inspected and flagged for removal if deemed artifactual.

The artifactual components were then subtracted from epoched data derived from the 0.10 Hz high pass filtered continuous data. Channels marked as bad were then interpolated using spherical splines, re-referenced to an average reference (recovering the FCz channel), and baseline corrected using the 200 ms immediately preceding stimulus onset. Finally, epochs were inspected for artifacts using the same routine described above. On average, 4.55% (range: 0.00-16.67%) of epochs in young adults and 4.14% (range: 0.31-15.74%) of epochs in older adults were rejected.

### ERP Measures

ERPs for scene, object, and face trials were created by averaging the artifact free epochs for the first presentation trials. The number of artifact free epochs for first presentation scene, object, and face trials were, respectively, 91 (range: 76-96), 92 (range: 81-96), and 91 (range: 80-96) for young adults and 92 (range: 77-96), 92 (range: 81-96), and 92 (range: 80-96) for older adults.

The ERPs were trimmed in time between -0.2 secs to 0.6 secs and low-pass filtered at 20 Hz with a finite impulse response filter (zero-phase shift; -6 dB cutoff = 22.5 Hz; transition bandwidth = 5 Hz) to reduce the bias in peak amplitude measures caused by high frequency noise (Kappenman et al., 2021; Luck, 2014). The ERP analysis focused on measuring the N170 and P200 from homotopic electrode clusters in the left (average of PO7 and P7) and right (average of PO8 and P8) hemispheres. These electrodes were selected for this analysis based on previous research demonstrating that the N170 and P200 effects are largest over lateral parieto-occipital channels (e.g., Boutet et al., 2020; Daniel & Bentin, 2012; R. Gao, 2015; Harel et al., 2016; Kappenman et al., 2021).

The ERP measures estimated for the analyses reported below included peak amplitude, peak latency, and 50% fractional peak onset for the P200 and N170. These measures were calculated from difference waveforms between scenes and objects for the P200, and between faces and objects for the N170. All measures were based on finding a local peak in the search window (0.125-0.250 sec time window for the scene-selective P200 and 0.090-0.170 sec for the face-selective N170). These windows were selected based on prior research of the P200 and N170 (e.g., Boutet et al., 2020; Daniel & Bentin, 2012; R. Gao, 2015; Harel et al., 2016; Kappenman et al., 2021). The windows were slightly wider than previous studies to accommodate potential age-related delays in peak amplitudes. Note that computation of the 50% fractional peak onset can occur below the lower-bound of the time window used for identifying the peak amplitude.

### Measurement of Neural Noise: 1/f aperiodic signal

We estimated neural noise from the prestimulus time window by quantifying the 1/f aperiodic component of the EEG power spectrum. The power spectrum was computed using Welch’s (1967) method on each epoch’s prestimulus window (200 samples per window with 75% overlap; zero-padded). The power spectrum for each channel was estimated by taking the across-epoch median power for frequencies in the 2-30 Hz range, and was then submitted to the FOOOF toolbox (Donoghue et al., 2020). This toolbox uses an algorithm that decomposes each channel’s power spectrum into oscillatory peaks and the aperiodic component. The aperiodic exponent was extracted from the 2-30 Hz frequency range of each channel’s power spectrum (*peak_width_limits = [2*.*0, 12*.*0], aperiodic_mode = ‘fixed’, peak_threshold = 1*). For each participant, we extracted the 1/f exponent from occipital and parietal electrodes (i.e., all O, PO, and P electrodes) given that our primary ERP effects of interest occur over posterior channels.

### Experimental Design and Statistical Analysis

Statistical analyses were conducted in *R* (version 4.1.1; R Core Team, 2021) with the following packages: *afex* (version 1.0; Singmann et al., 2021), *modelbased* (version 0.8.1; Makowski et al., 2020), and *effectsize* (version 0.4.5, Ben-Shachar et al., 2020). When necessary, the Holm (1979) procedure was used to correct for multiple comparisons. Results are considered significant at a two-tailed *p* < .05 unless otherwise specified.

#### Neuropsychological Test Performance

Age differences in neuropsychological test performance were examined with independent samples *t*-tests assuming equal variance using the *t*.*test* function in *R*.

#### Behavioral Performance on the 1-Back Task

The RTs for repeated (i.e., 1-back) trials receiving a correct response were submitted to a generalized linear mixed effects model with an inverse Gaussian link function (Lo & Andrews, 2015) using the *mixed* function of the *afex* package. This model included fixed effects of age group (young, older), image category (scene, object, and face) and the age group by image category interaction. The model included a maximal random effects structure with a random intercept for participant and a random slope for image category, as well as the covariance between the random effects. Note that the null hypothesis significance test for the fixed effects were based on likelihood ratio tests, and post-hoc contrasts conducted with the *modelbased* package were based on asymptotic (i.e., *z*) tests.

#### ERP Peak Measures

The ERP peak amplitude, latency, and fractional peak onset measures were examined using a linear mixed effects model using the *mixed* function in the *afex* package. The model had fixed effects of age group (young, older), hemisphere (left, right), and the age group by hemisphere interaction and a random intercept term. Post-hoc contrasts on the interaction, when necessary, were based on *t*-tests conducted using the *modelbased* package. For these linear mixed effects models, and those reported below, degrees of freedom used the Satterthwaite (1946) approximation.

To control for individual differences in vision, we reran the above models including mean centered visual acuity (logMAR) and contrast acuity (logCS). With one exception, inclusion of these covariates did not affect the pattern of findings (also see Koen et al., 2019). We report the results from the models that do not include these covariates for the sake of parsimony. The one exception is noted in the Results (see the Face N170 subsection of ERP Peak Measures).

#### 1/f Aperiodic Signal

Age differences in neural noise were investigated by submitting the 1/f aperiodic component (i.e., exponent) of the prestimulus power spectrum from parietal-occipital channels to a linear mixed effects model. This model included age group as the only fixed effect factor.

Random intercepts for participant and channel were also included in this model.

We also investigated the relationship between neural noise (measured with the 1/f aperiodic exponent) and the ERP peak measures for the P200 and N170 using linear mixed effects models. For this analysis, a participant specific neural noise measure was created by averaging over all the 1/f exponents estimated from the posterior electrodes (i.e., P, PO, and O). This model included fixed effects of age group (young, older), hemisphere (left, right), 1/f exponent, and the interactions between all three factors, along with a random intercept. Note that the 1/f exponent was mean centered prior to estimating the model. In these models, we focused only on effects involving the 1/f exponent measure. Post-hoc contrasts on interactions involving the 1/f exponent were conducted by testing if the simple slopes within each factor (or combination of factors) was significantly different from 0 using the *modelbased* package.

As with the ERP peak measures, we included mean centered visual and contrast acuity as covariates to control for individual differences in visual function. Inclusion of these covariates did not alter the pattern of findings (also see Koen et al., 2019). Thus, we report the results from the statistical models that do not include these covariates.

## Results

### Neuropsychological Test Performance

Table 1 provides the sample demographics and performance on the neuropsychological test battery. Older adults had significantly more education than young adults. Statistically significant age-related reductions were observed in every measure, except for the Vocabulary subtest of the WASI-II and the TOPF which showed no age differences.

### Behavioral Results

Participants demonstrated high accuracy on detecting the repeats (young adults: M = 0.96, SE = 0.01; older adults: M = 0.95, SE = 0.01) and correctly refrained from responding on first presentation trials (young adults: M = 0.999, SE = 0.000; older adults: M = 0.997, SE = 0.001).

The generalized linear mixed effects model on trial-wise reaction time data revealed a significant effect of age group [χ^2^(1) = 7.46, *p* = .006]. As expected, older adults were slower to respond to repeated images relative to younger adults (see Figure 1B). There was also a significant effect of image category [χ^2^(2) = 14.69, *p* = .001]. Post-hoc contrasts demonstrated that reaction times to repeated scenes were significantly slower compared to both repeated faces [*z* = 3.20, *p* = .004, Cohen’s *d* = 0.34] and repeated objects [*z* = 3.69, *p* = .001, Cohen’s *d* = 0.39]. There was no significant difference in reaction times for correctly identifying repeated faces and objects [*z* = 0.29, *p* = .956, Cohen’s *d* = 0.03] (see Figure 1C). The age group by image category interaction was not significant [χ^2^(2) = 2.24, *p* = .326].

### ERP Peak Measures

The grand average waveforms from the three image categories in young and older adults are shown in Figures 2 and 3, respectively. Visual inspection of these waveforms in both age groups show differences between ERPs for scene and object stimuli around 0.15 to 0.25 secs following stimulus onset. These differences are consistent with the scene-selective P200 previously reported by Harel and colleagues (2016). Similarly, consistent with prior work on the face N170 (Boutet et al., 2020; Daniel & Bentin, 2012; L. Gao et al., 2009; Kappenman et al., 2021; Rousselet et al., 2007, 2009; for review, see Rossion & Jacques, 2012), both young and older adults showed a more negative going ERP for faces relative to objects between 0.1 and 0.17 secs following stimulus onset. Difference waveforms were used to measure the peak amplitudes, peak latencies, and fractional peak onsets for the P200 (scene minus object) and N170 (face minus object).

**Figure 2.**
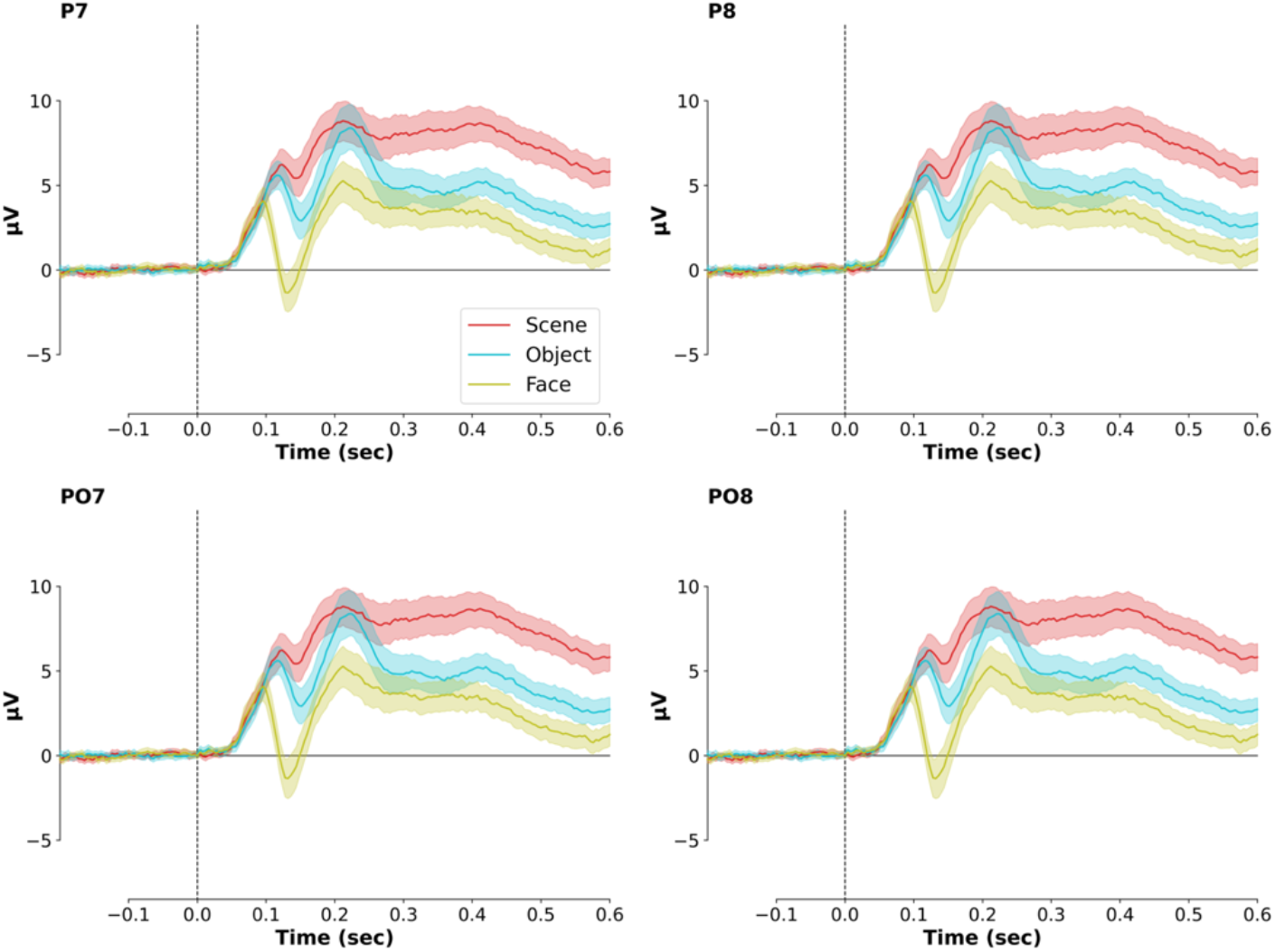
Grand average ERPs for young adults for scenes, objects, and faces. The electrodes shown were those used to create virtual left (P7 and PO7) and right (P8 and PO8) hemisphere channels for analysis of the P200 and N170 in the main text. The ERPs reflect the grand average of unfiltered ERPs for each image category and error ribbon reflects the 95% confidence interval.

**Figure 3.**
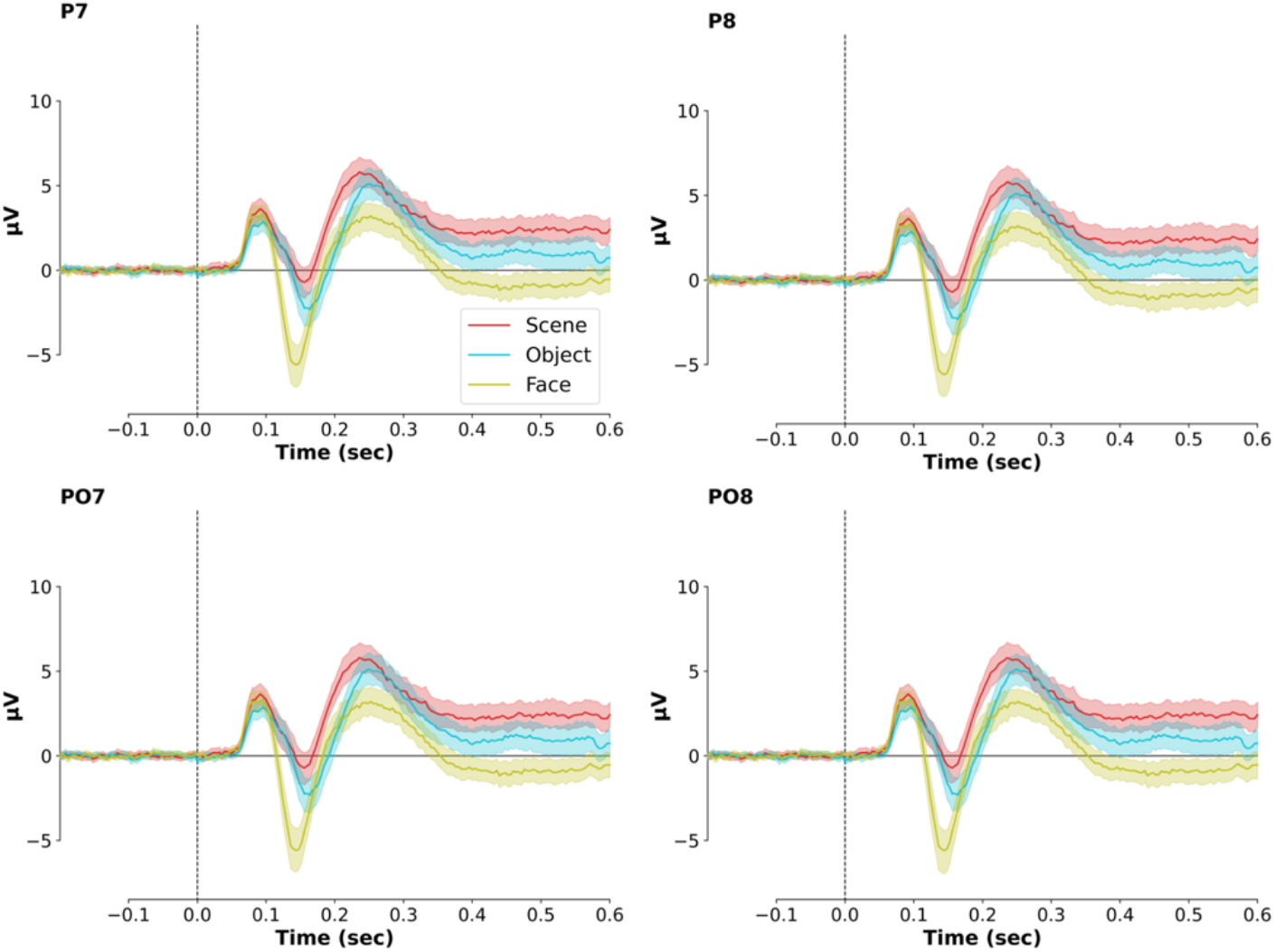
Grand average ERPs for older adults for scenes, objects, and faces. The electrodes shown were those used to create virtual left (P7 and PO7) and right (P8 and PO8) hemisphere channels for analysis of the P200 and N170 in the main text. The ERPs reflect the grand average of unfiltered ERPs for each image category and error ribbon reflects the 95% confidence interval.

#### Scene P200

Figure 4A shows the grand average difference waveforms between scene and object ERPs for young and older adults, and Table 2 includes the descriptive statistics for P200 peak measures. A visual inspection of the waveforms indicates that the peak amplitude in the 0.15 to 0.25 second time window was larger and occurred earlier (in onset and peak latency) in young relative to older adults. The results from the analysis of the P200 peak measures support these observations. The linear mixed effects model for the scene selective P200 peak amplitude (Figure 4B) showed a significant main effect of age group [*F*(1, 86) = 15.01, *p* < .001, 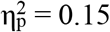], hemisphere [*F*(1, 86) = 4.44, *p* = .038, 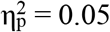], and the age group by hemisphere interaction [*F*(1, 86) = 5.04, *p* = .027, 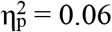]. The main effect of age group was driven by the peak amplitude of the P200 being reduced in magnitude for older relative to young adults (see Table 2). The post-hoc analysis of the interaction indicated that the above age-related reduction in P200 peak amplitude was significant in the right hemisphere [*t*(154.47) = 4.46, *SE* = 0.40, *p* < .001, Cohen’s *d* = 0.72]. However, the age difference in the left hemisphere only approached, but did not reach, our *a priori* significance threshold [*t*(154.47) = 1.88, *SE* = 0.40, *p* = .063, Cohen’s *d* = 0.30]. Overall, these findings converge with previous fMRI findings of age-related neural dedifferentiation of scene stimuli in the parahippocampal place area (Koen et al., 2019; D. C. Park et al., 2004; Srokova et al., 2020; Voss et al., 2008).

**Table 2.** P200 peak amplitudes and latencies means and standard error for age group and hemispheres.

| Age Group | Hemisphere | Peak Amplitude | Peak Latency | Fractional Peak Onset Latency |
| --- | --- | --- | --- | --- |
| Young | Left | 4.68 (0.27) | 0.169 (0.002) | 0.136 (0.003) |
|  | Right | 5.68 (0.31) | 0.165 (0.002) | 0.130 (0.004) |
| Older | Left | 3.94 (0.28) | 0.193 (0.003) | 0.160 (0.004) |
|  | Right | 3.91 (0.25) | 0.196 (0.002) | 0.162 (0.003) |

*Note.* Standard error of the mean is provided in parentheses. Peak amplitude is reported in $\mu V$ and latency measures are reported in secs.

**Figure 4.**
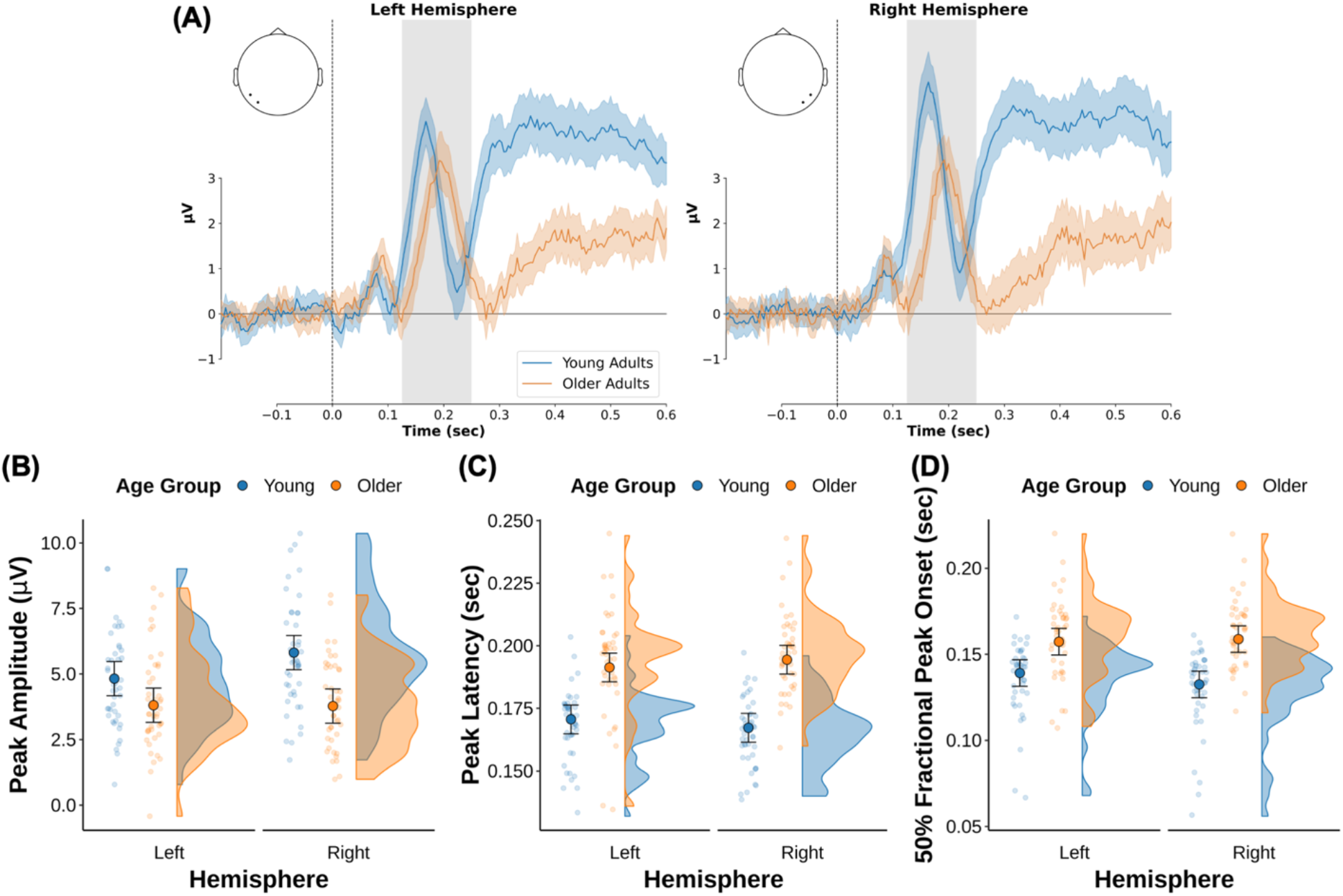
Results from the analysis of the scene selective P200. (A) Grand average difference waveforms between scene and object ERPs from virtual channels from the left (average of P7 and PO7) and right (average of P8 and PO8) hemispheres. These waveforms reflect the grand averages of unfiltered ERPs from young and older participants, with the error ribbon reflecting the 95% confidence interval of the difference wave in each age group. The shaded area reflects the time window used to search for the P200 peak amplitude measure. Note that the peak measures were derived from ERPs that were low-pass filtered with a 20 Hz finite impulse response filter (zero-phase shift; -6 dB cutoff = 22.5 Hz; transition bandwidth = 5 Hz). (B-D) Raincloud plots of the P200 peak amplitude, latency, and 50% fractional peak onset latency derived from the scene minus object difference wave. The peak amplitude of the P200 was larger in young relative to older adults and only significant in the right hemisphere. Both P200 peak latency and fractional onset were delayed in older relative to young adults. The point with the error bar represents the group mean and 95% confidence interval.

The linear mixed effects model conducted on the P200 peak latency revealed a main effect of age group [*F*(1, 86) = 96.49, *p* < .001, 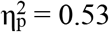] that was driven by the peak P200 latency in older adults occurring later relative to young adults (Figure 4C and Table 2). Neither the main effect of hemisphere [*F*(1, 86) = 0.00, *p* = .946, 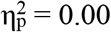] nor the age group by hemisphere interaction [*F*(1, 86) = 2.54, *p* = .114, 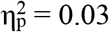] reached our *a priori* statistical threshold.

The linear mixed effects model conducted on the P200 fractional peak onset latency revealed a main effect of age group [*F*(1, 86) = 49.32, *p* < .001, 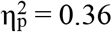] that was driven by the P200 fractional peak onset latency in older adults occurring later relative to young adults (Figure 4D and Table 2). Neither the main effect of hemisphere [*F*(1, 86) = 1.17, *p* = .282, 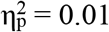] nor the age group by hemisphere interaction [*F*(1, 86) = 3.02, *p* = .086, 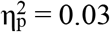] reached our *a priori* statistical threshold. The peak latency and onset findings suggest that, in addition to agerelated reductions in the magnitude of neural selectivity, there is also an age-related delay in when neural activity shows selectivity for scene stimuli. Inclusion of visual and contrast acuity measures as covariates did not change the pattern of results for the above analyses (see Experimental Design and Statistical Analyses).

#### Face N170

Figure 5A shows the grand average difference waveforms between face and object ERPs for young and older adults, and Table 3 includes the descriptive statistics for N170 peak measures. Visual inspection of these difference waveforms suggests that the N170 was larger (i.e., more negative) and delayed in onset and latency in young relative to older adults. A linear mixed effects model conducted on the N170 peak amplitude (Figure 5B) revealed a main effect of age group [*F*(1, 86) = 5.43, *p* = .022, 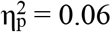] that was driven by older adults having smaller (i.e., less negative) peak N170 amplitudes compared to young adults (see Table 3). This finding is consistent with fMRI studies of age-related neural dedifferentiation in the fusiform face area (D. C. Park et al., 2004; J. Park et al., 2012; Voss et al., 2008; but see Srokova et al., 2020).

**Table 3.** N170 peak amplitudes and latencies means and standard error for age group and hemispheres.

| Age Group | Hemisphere | Peak Amplitude | Peak Latency | Fractional Peak Onset Latency |
| --- | --- | --- | --- | --- |
| Young | Left | -7.27 (0.52) | 0.128 (0.002) | 0.109 (0.002) |
|  | Right | -7.18 (0.65) | 0.127 (0.002) | 0.106 (0.002) |
| Older | Left | -6.14 (0.44) | 0.137 (0.002) | 0.118 (0.002) |
|  | Right | -5.45 (0.36) | 0.138 (0.002) | 0.120 (0.002) |

*Note.* Standard error is provided in parentheses with the mean values. Peak amplitude is reported in $\mu V$ and latency measures are reported in secs.

**Figure 5.**
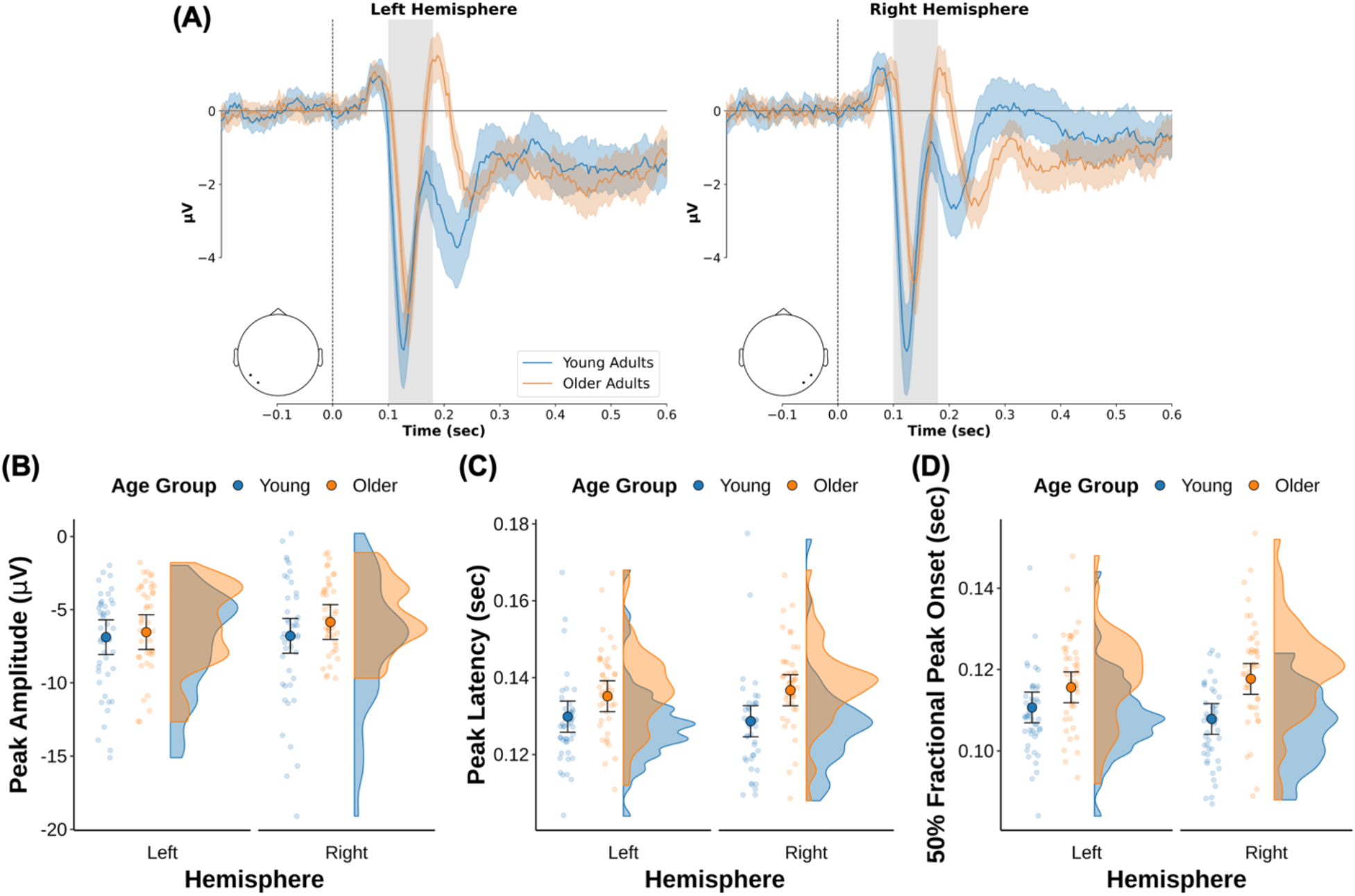
Results from the analysis of the face selective N170. (A) Grand average difference waveforms between scene and object ERPs from virtual channels from the left (average of P7 and PO7) and right (average of P8 and PO8) hemispheres. These waveforms reflect the grand averages of unfiltered ERPs from young and older participants, with the error ribbon reflecting the 95% confidence interval of the difference wave in each age group. The shaded area reflects the time window used to search for the N170 peak amplitude measure. Note that the peak measures were derived from ERPs that were low-pass filtered with a 20 Hz finite impulse response filter (zero-phase shift; -6 dB cutoff = 22.5 Hz; transition bandwidth = 5 Hz). (B-D) Raincloud plots of the N170 peak amplitude, latency, and 50% peak onset latency derived from the scene minus object difference wave. The peak amplitude of the N170 was larger in young relative to older adults, and the N170 peak latency and fractional peak onset were delayed in older adults relative to young adults. The point with the error bar represents the group mean and 95% confidence interval.

Neither the main effect of hemisphere [*F*(1, 86) = 1.19, *p* = .279, 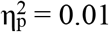] nor the age group by hemisphere interaction [*F*(1, 86) = 0.70, *p* = .406, 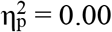] reached our *a priori* statistical threshold. However, the age difference in N170 amplitude was no longer significant visual and contrast acuity measures were included as covariates [*F*(1, 84) = 0.51, *p* = .476, 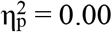]. This finding suggests that, unlike the scene P200, age differences in the N170 were not independent of individual differences in visual function.

A similar linear mixed effects model on the N170 peak latency revealed a main effect of age group [*F*(1, 86) = 21.77, *p* < .001, 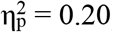] that was driven by the peak N170 latency in older adults occurring later relative to young adults (Figure 5C and Table 3). Neither the main effect of hemisphere [*F*(1, 86) = 0.02, *p* = .881, 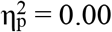] nor the age group by hemisphere interaction [*F*(1, 86) = 1.26, *p* = .265, 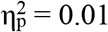] reached our *a priori* statistical threshold.

A linear mixed effects model conducted on the N170 fractional peak onset latency revealed a main effect of age group [*F*(1, 86) = 32.90, *p* < .001, 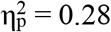] that was driven by the N170 fractional peak onset latency in older adults occurring later relative to young adults.

Neither the main effect of hemisphere [*F*(1, 86) = 0.09, *p* = .770, 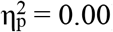] nor the age group by hemisphere interaction [*F*(1, 86) = 3.94, *p* = .050, 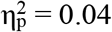] reached our *a priori* statistical threshold. The interaction did, however, approach our significance threshold, which is likely due to larger age differences in fractional onset latency in the right versus left hemisphere (see figure 5D and Table 3). The peak latency and onset findings suggest that there are also age-related delays in the face selective N170 (for similar findings, see Boutet et al., 2020; Daniel & Bentin, 2012; but see L. Gao et al., 2009). Note that inclusion of visual and contrast acuity did not alter the latency or onset results.

### Age Differences in Neural Noise (1/f Exponent)

The data from the analysis of the 1/f exponent is shown in Figure 6. The exponents derived from posterior channels were analyzed for age differences with a linear mixed effects model that included random intercepts for participant and channel (see Experimental Design and Statistical Analyses). There was a significant age-related reduction in the 1/f exponent [*F*(1, 86) = 52.97, *p* < .001, 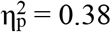]. This finding is consistent with results from prior studies examining age-related changes in aperiodic electrophysiological activity (Dave et al., 2018; Voytek et al., 2015; also see Voytek & Knight, 2015) and suggests that neural noise is greater in older relative to young adults (cf. R. Gao, 2015).

**Figure 6.**
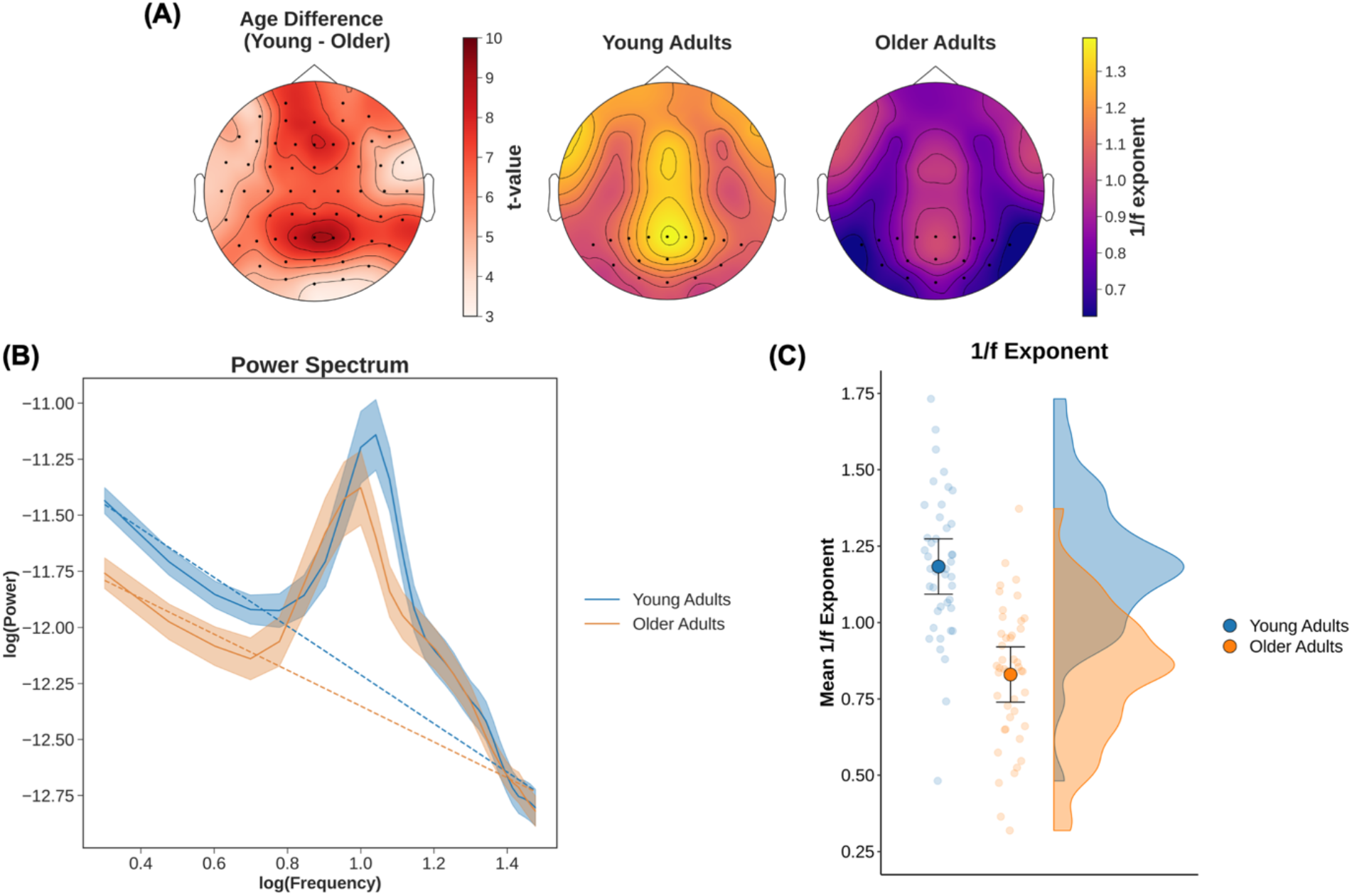
Results from the analysis of age differences in the 1/f exponent. The 1/f exponent was estimated by applying the FOOOF algorithm (Donoghue et al., 2020) to the power spectrum derived from the 1 sec pre-stimulus baseline window. (A) The left panel shows the topography of the age differences in the 1/f exponent. Positive *t*-values reflect larger 1/f exponents in young relative to older adults. The right panel shows the topography of the across-participant average 1/f exponent estimates for young and older adults. The black points in the right panel reflect the posterior electrodes (P, PO, and O electrodes) used for the analyses depicted in (B) and (C). (B) The average power spectrum across the posterior channels for young and older adults plotted in log-log space. The error ribbon depicts the 95% confidence interval of the estimate. The estimate 1/f exponent (i.e., slope) estimated from the grand average power spectrum is shown as the dotted line. (C) Participant estimates of the 1/f exponent averaged across the posterior electrodes shown in (A). The point with the error bar represents the group mean and 95% confidence interval. Age differences were largest over central posterior electrodes. Larger 1/f exponents (i.e., steeper slopes) reflect less aperiodic activity in the power spectrum and indicate lower levels of neural noise. Thus, the reduction of the estimate 1/f exponents (i.e., shallower slopes) in older adults suggest age-related increases in neural noise.

### Relationship Between Neural Selectivity and Neural Noise

Computational models of neural dedifferentiation propose that increased neural noise is related to reductions in neural selectivity (Li et al., 2001; Li & Rieckmann, 2014a). We tested this prediction by examining the association between the ERP peak measures (i.e., amplitude, latency, and onset latency) and the average of the 1/f exponents derived from the posterior channels used in the above analysis. The linear mixed effects models reported in this section are like the statistical models reported on in the Scene P200 and Face N170 sections above, but they include the 1/f exponent (mean centered) as a predictor along with the interaction between the other terms in the model (i.e., age group by 1/f interaction, hemisphere by 1/f interaction, and the 3-way interaction between age group, hemisphere, and 1/exponent). In this section we focus only on effects that included the 1/f exponent. Inclusion of visual and contrast acuity measures as covariates did not change the pattern of results for the above analyses (see Experimental Design and Statistical Analyses).

#### P200

The linear mixed effects model for the scene selective P200 peak amplitude revealed a significant age group by 1/f exponent interaction [*F*(1, 84) = 7.53, *p* = .007, 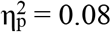]. This interaction was driven by a significantly steeper slope in older adults relative to younger adults (Figure 7). Post-hoc analyses demonstrated that there was a negative relationship between the 1/f exponent and the P200 amplitude in older adults that was significantly different from zero [*slope* = 2.75, *t*(84) = 2.75, *p* = .007]. This relationship demonstrates that older adults with low levels of neural noise showed higher P200 peak amplitudes. The relationship between the 1/f exponent and P200 peak amplitude was not significant in young adults [*slope* = -1.06, *t*(84) = 1.10, *p* = .274]. No other effects involving the 1/f exponent were significant [all *F*’s < 1.49, all *p*’s > .226, all 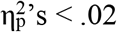].

**Figure 7.**
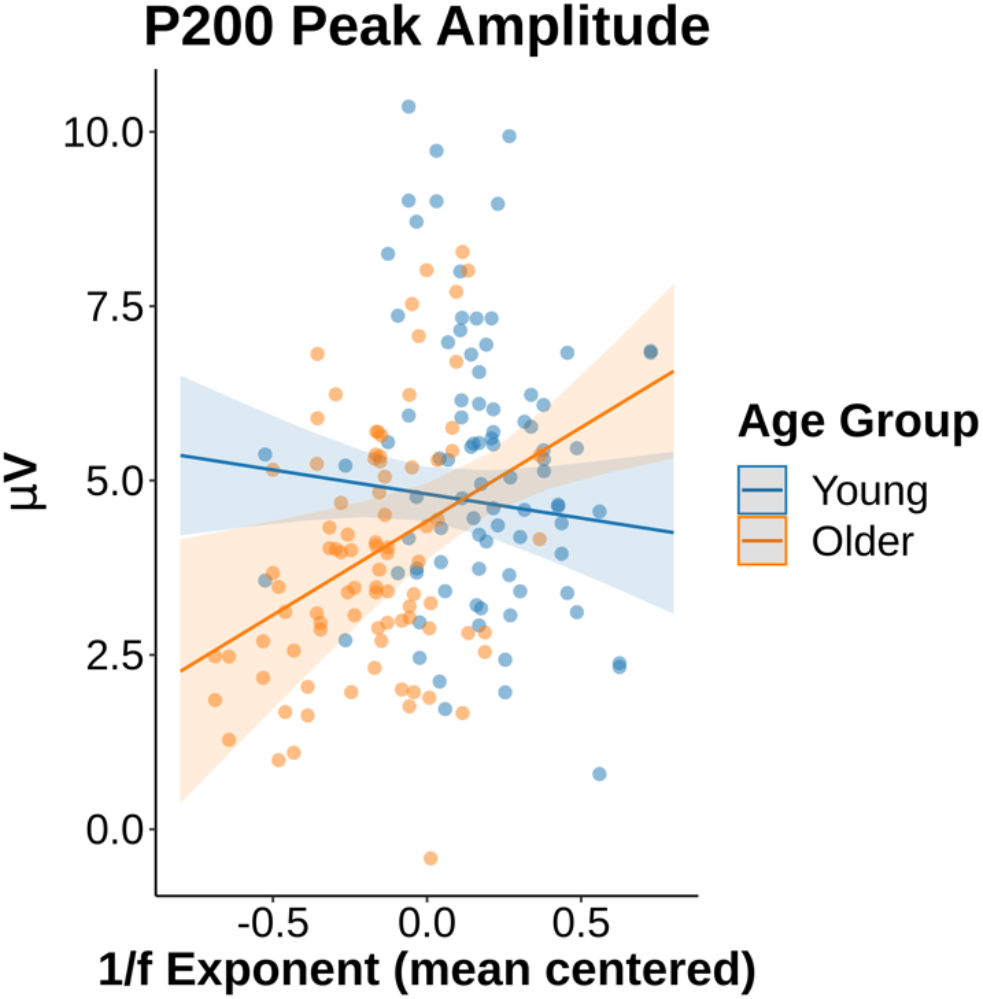
Visual depiction of the age group by 1/f exponent interaction for the analysis of the P200 peak amplitude data. There was a significant relationship between the 1/f exponent and the P200 peak amplitude in older, but not younger, adults. In older adults, lower levels of neural noise (i.e., higher 1/f exponents) were associated with higher P200 peak amplitudes. The error bar around the regression reflects the 95% confidence interval and the circles represent individual data points.

The linear mixed effects model for the scene selective P200 peak latency revealed a significant interaction between age group, hemisphere, and 1/f exponent [*F*(1, 84) = 3.98, *p* = .049, 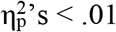]. To examine the interaction, we conducted post-hoc analyses on the four simple slopes for each hemisphere in each age group. After correction for four comparisons using the Holm method, none of the simple slopes were significantly different from 0 [all slopes between - .013 and .015; all *t*(149.34) < 1.30, all *p*Holm > .780]. No other effects involving the 1/f exponent were significant [all *F*’s < 1.26, all *p*’s > .266, all 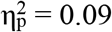].

The analysis of the P200 fractional peak onset showed a significant 3-way interaction involving age group, hemisphere, and 1/f exponent [*F*(1, 84) = 9.31, *p* = .003, 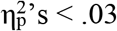].

However, similar to P200 peak latency, none of the simple slopes were significant after correction for multiple comparisons [all slopes between -.032 and .016; all *t*(130.78) < 2.16, all *p*Holm > .129]. No other effects involving the 1/f exponent were significant [all *F*’s < 3.31, all *p*’s > .071, all 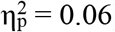]. Together, these latter two findings suggest that, at best, the 1/f exponent measure of neural noise has a rather weak (and likely complex) relationship with the P200 peak latency and fractional peak onset.

#### N170

The linear mixed effects model for the face selective N170 peak amplitude showed a significant main effect of 1/f exponent [*F*(1, 84) = 5.09, *p* = .027, 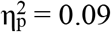] that was moderated by an interaction with hemisphere interaction [*F*(1, 84) = 8.38, *p* = .005, 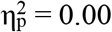] (see Figure 8). An analysis of the simple slopes indicate a significant negative relationship between N170 amplitude and the 1/f exponent in the left hemisphere [*slope* = -5.25, *t*(133.83) = 3.40, *p* < .001] but not in the right hemisphere [*slope* = -0.80, *t*(133.83) = 0.52, *p* = .605]. Thus, in the left hemisphere, there was an age-invariant relationship showing 1/f exponents (increases in neural noise) are associated with decreased neural selectivity. Neither the age group by 1/f exponent interaction [*F*(1, 84) = 0.05, *p* = .815, 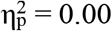] nor the age group by 1/f exponent by hemisphere interaction [*F*(1, 84) = 0.22, *p* = .637, 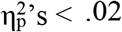] reached our *a priori* statistical threshold.

**Figure 8.**
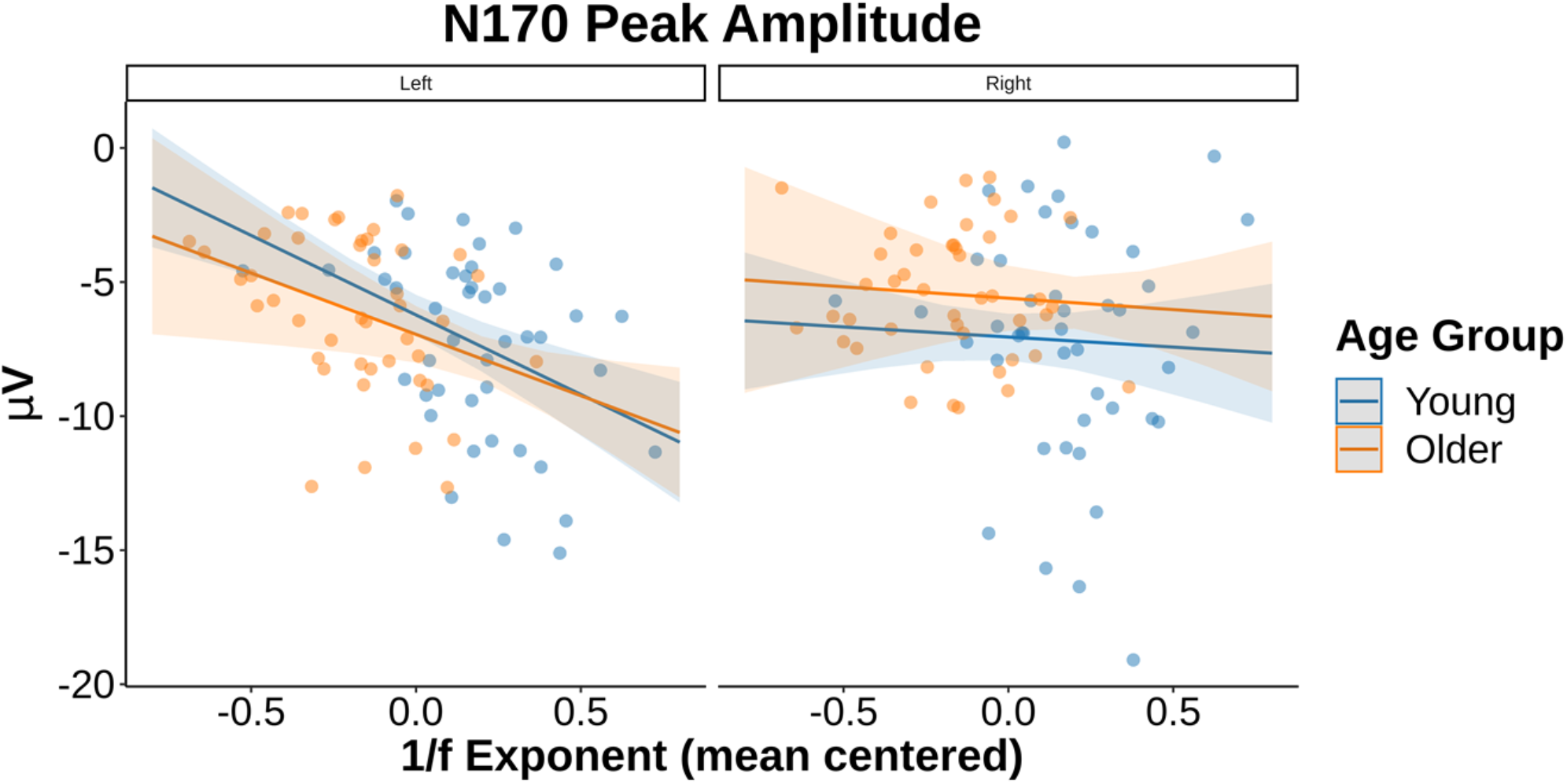
Results from the analysis of the face selective N170 and 1/f exponent. Data points for the N170 peak amplitudes and 1/f exponents for young and older adults are presented, with slopes representing group mean estimates and error ribbons reflecting the 95% confidence interval of group mean estimates from the linear mixed effects model for each age group. In the left hemisphere, more negative 1/f exponents reflecting less aperiodic activity in the power spectrum and indicating less neural noise are associated with smaller N170 peak amplitudes in both young and older adults. Thus, in the left hemisphere, the age-related reduction of 1/f exponents (increases in neural noise) are associated with decreased neural selectivity in an age-invariant manner. In other words, more lateralized N170 peak amplitudes are associated with decreased neural noise in an age-invariant manner.

None of the effects that included the 1/f exponent reached our significance threshold for the linear mixed effects models of N170 peak latency [all *F*’s < 2.90, all *p*’s > .092, all 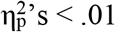] nor fractional peak onset [all *F*’s < 1.25, all *p*’s > .268, all 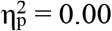].

### General Discussion

The current study examined the relationship between neural noise and age differences in EEG measures of neural selectivity (Li et al., 2001; Li & Rieckmann, 2014b). Specifically, we related measures of neural noise derived from the aperiodic activity in the EEG signal (Dave et al., 2018; Donoghue et al., 2020; R. Gao, 2015; Voytek et al., 2015) and ERP measures of neural selectivity, namely the scene-selective P200 (Harel et al., 2016) and face-selective N170 (for review, see Rossion & Jacques, 2012). First, the results demonstrated that the amplitude of the scene-selective P200, but not the face-selective N170, was reduced in older adults relative to younger adults (after controlling for individual differences in visual and contrast acuity). Second, both the P200 and N170 components showed evidence of age-related slowing. Third, in older but not young adults, the P200 peak amplitude was associated with the 1/f exponent, indicating that lower levels of neural noise were associated with higher P200 peak amplitudes, or neural selectivity. Finally, the N170 peak amplitude in the left, but not right, hemisphere showed an age-invariant relationship with the 1/f measure of neural noise whereby lower levels of neural noise were associated with larger (i.e., more negative) N170 amplitudes.

### Age Differences in ERP Neural Selectivity Measures

The present findings replicate those from a recent study by Harel and colleagues (2016), who provided the first demonstration of this scene-selective P200 ERP component. Moreover, we extended these findings to show that the scene-selective P200 is observed in both young and older adults, suggesting that this scene-selective component can be generalized across different groups of individuals.

Our findings also provide the first demonstration of age-related neural dedifferentiation using ERPs in paradigms like those used in fMRI research (e.g., D. C. Park et al., 2004; Voss et al., 2008). These findings converge with fMRI studies demonstrating that neural dedifferentiation is more readily observed for scene stimuli relative to face stimuli (Koen et al., 2019; Srokova et al., 2020; for related findings, see Voss et al., 2008). This is supported by our results showing age-related reduction in the amplitude of the scene-selective P200, but not the face-selective N170 (after controlling for visual and contrast acuity). These findings suggest that there is something special about scene stimuli that produces consistent patterns of age-related neural dedifferentiation.

The present results also extend prior fMRI findings in showing that age differences in neural selectivity reflect a mixture of reductions in amplitude of and delays in neural processing. For both the P200 and the N170, we observe significant age-related delays in onset and peak latency, which suggests that the neural processes that differentiate categories of visual stimuli are sensitive to age-related slowing. It is possible that mixed findings for age-related neural dedifferentiation for faces (Srokova et al., 2020; and potentially objects; see Koen et al., 2019) observed with fMRI are driven by age-related delays in the peak level of neural selectivity, but not overall differences in the magnitude of neural selectivity. Future research using a multi-model approach in the same participants is needed to critically examine this hypothesis and to determine if the present EEG findings for scenes reflect similar neural processes as those observed in fMRI studies of neural dedifferentiation.

Our results also add to the existing literature on age differences in the latency of the face-selective N170. Previous studies have reported mixed findings with some showing prolonged peak latencies in older relative to young adults (Boutet et al., 2020; Daniel & Bentin, 2012; Rousselet et al., 2009), whereas others have reported no age differences in latency (L. Gao et al., 2009). Our results converge with the former set of findings, suggesting there are age-related delays in the N170. Importantly, our methods differ from prior investigations that focused on peak latency measures from ERPs for face stimuli alone. Here, we measured the peak latency (as well as fractional peak onset) from difference waves. We believe this approach can better isolate underlying neural components given that peaks are not (necessarily) the same as components (Luck, 2014). Moreover, our different measurement method could potentially explain why the N170 amplitude results differed from the prior N170 literature.

It is unclear from the present results if age differences in the two ERP components investigated are driven by age differences in neural activity elicited by scenes and faces, or a common age difference in the neural response to just the object stimuli. Specifically, both the N170 and P200 were estimated from difference waves created by referencing the ERPs elicited by faces and scenes to the ERPs elicited by object stimuli. Thus, it is possible that age differences in object ERPs could account for the observed pattern of results. Indeed, age differences in the ERPs elicited by objects can be seen by comparing Figure 2 and 3. However, it should be noted that even age differences in ERPs for a single condition (e.g., ERP for object stimuli) can occur even when young and older adults have the same underlying neural generator. Such differences can arise from age-related differences in neuroanatomy, which can alter the orientation or generator-to-scalp distance of a neural generator, thus affecting the electrical activity recorded at the scalp for any single-condition ERPs. The use of difference waves helps to mitigate this issue. Although we cannot fully rule out the possibility that age differences in the object ERP contribute to the present results, we believe this account unlikely as it cannot readily explain the different patterns observed for the P200 and N170 measured from difference waves.

### Relationship Between Age, Neural Noise, and Neural Selectivity

Another aim of this study was to examine the relationship between neural noise and neural dedifferentiation in young and older adults. Computational models propose that agerelated increases in neural noise contribute to age-related reductions in neural selectivity (Li et al., 2001; Li & Rieckmann, 2014a). In addition to replicating previous findings showing age-related increases in neural noise as indexed by decreases in the 1/f exponent (Dave et al., 2018; Voytek et al., 2015), our results provide some evidence consistent with the general proposal that neural noise is associated with neural selectivity and, more specifically, the magnitude (not latency) of neural selectivity. Importantly, the relationship between neural noise and neural selectivity appeared to differ by age and visual category.

For the scene-selective P200, we found the relationship between neural noise and peak amplitude was moderated by age group. In older adults, lower levels of neural noise (i.e., higher 1/f exponents) were associated with higher levels of neural selectivity for scene stimuli (i.e., larger P200 peak amplitudes). There was no significant relationship between neural noise and the P200 peak amplitude in younger adults. This pattern of results might suggest a brain maintenance or preservation account of age-related neural dedifferentiation (Cabeza et al., 2018). Specifically, it could be the case that older adults who have maintained levels of neural noise more similar to those found in young adults through some lifestyle factors, such as exercise (Kleemeyer et al., 2017), maintain young adult levels of neural selectivity. A longitudinal study of this relationship is needed to remove potential cohort effects and provide better evidence for a the brain maintenance account mentioned above.

In contrast, peak amplitude of the face-selective N170 showed an age-invariant relationship with neural noise that was only observed in left hemisphere electrodes. While the age-invariant nature of this relationship is in line with predictions derived from computational models (Li et al., 2001), this observed laterality effect was unexpected. One possible interpretation of this finding is that the right lateralization typically reported in the N170 literature is dependent on levels of neural noise. Specifically, both young and older adults with lower levels of neural noise also show reduced hemispheric asymmetry in the N170 compared to individuals with higher levels of neural noise. These findings need to be replicated before any strong conclusions can be drawn made about this hemispheric dependent, yet age-invariant, relationship between neural noise and the N170 peak amplitude.

### Implications for Computational Models of Neural Dedifferentiation Theory

While our results are broadly consistent with the predicted relationship between neural noise and neural selectivity derived from computational models, the present results join a growing body of work suggesting that computational models of neural dedifferentiation require revision (Koen et al., 2019; Srokova et al., 2020; but see, Simmonite & Polk, 2022). As we have argued previously, computational models of neural dedifferentiation predict age- and stimulus-invariant relationships between neural noise and measures of neural selectivity (Koen et al., 2020; Koen & Rugg, 2019). The present findings join the growing party of evidence that neural dedifferentiation is not consistently observed across all stimulus categories. Moreover, the present results suggest that the relationship between neural noise and neural dedifferentiation differs across the lifespan and, potentially, in different brain regions. The present results provide some important boundary conditions that any new iteration of a computational model of neural dedifferentiation must account for.

## Acknowledgements

This work was supported by an award to J.D.K. from the National Institute on Aging grant R56AG049583. We would like to thank Seham S. Kafafi, Sarah N. Berland, Sofia Zitella, Margaret Allen, and Lyndsey Della Vecchia for providing feedback on previous draft of this manuscript.

